# Load Non-Linearly Modulates Movement Reflex Gain in an Insect Leg via a Distributed Network of Identified Nonspiking Interneurons

**DOI:** 10.1101/2022.02.24.481822

**Authors:** Corinna Gebehart, Scott L. Hooper, Ansgar Büschges

## Abstract

Producing context-specific motor acts requires sensorimotor neural networks to integrate multiple sensory modalities. Some of this integration occurs via presynaptic interactions between proprioceptive afferent neurons themselves,^1,2^ other by afferents of different modalities targeting appropriate motor neurons (MNs).^3–5^ How the interneuronal network typically interposed between sensory afferents and MNs contributes to this integration, particularly at single-neuron resolution, is much less understood. In stick insects, this network contains nonspiking interneurons (NSIs) converging onto the posture-controlling slow extensor tibiae motor neuron (SETi). We analyzed how load altered movement signal processing by tracing the interaction of movement (femoral chordotonal organ, fCO) and load (tibial campaniform sensilla, tiCS) signals from the afferents through the NSI network to the motor output. On the afferent level, load reduced movement signal gain by presynaptic inhibition; tiCS stimulation elicited primary afferent depolarization and reduced fCO afferent action potential amplitude. In the NSI network, graded responses to movement and load inputs summed nonlinearly and increased the gain of NSIs opposing movement-induced reflexes. The gain of SETi and muscle movement reflex responses consequently decreased. Gain modulation was movement parameter-specific and required presynaptic inhibition; pharmacologically blocking presynaptic inhibition abolished load-dependent tuning of SETi responses. These data describe sensorimotor gain control at the sensory, premotor, and motor levels. Presynaptic inhibition-mediated nonlinear integration allowed the NSI network to respond to movement sensory input in a context (load)-dependent manner. These findings show how gain changes can allow premotor networks to integrate multiple sensory modalities and thus generate context-appropriate motor activity.

## INTRODUCTION

Neural networks process multiple input streams conveying qualitatively different information. In motor networks, these inputs include central drives, interactions with other motor networks, and sensory input monitoring internal (proprioception) and external (exteroception) state.^6,7^ In legged animals, premotor networks process these signals both during locomotion and while maintaining posture against gravity and other external perturbations.^5,8,9^

Maintaining functional motor output requires altering neural network response to sensory inputs in a context-specific manner. One mechanism to do so is global changes in synaptic strength. It is now apparent, however, that another important mechanism is differentially altering the gain of different input modalities^1^: abolishing gain control causes motor oscillations in goal-directed reaching in mice^10^ and gain control helps stabilize motor neuron (MN) activity, gate self-generated feedback, increase motor control precison,^11,12^ and optimize network performance.^12^ This sensitivity to gain arises because gain determines the *rate* of change of the system to changing input, which can be much more destabilizing than simple global changes in synaptic strength; in dynamics systems jargon, it makes the system more “brittle”. However, what constitutes optimization depends on the context in which the motor output occurs. Gain control therefore needs to be context-dependent, and presynaptic modulation of afferent input onto premotor networks, in both vertebrates and insects, does depend on behavioral context, e.g., locomotion.^13,14^

Motor networks in the spine (vertebrates) or ventral nerve cord (insects) receive proprioceptive information about joint movement and angle from muscle spindles^9,15–18^ (vertebrates) and chordotonal organs^7,19,20^ (insects) and forces acting on the limbs from Golgi tendon organs^9,15,17,18,21^ (vertebrates) and campaniform sensilla^7,20,22^ (CS, invertebrates). Studying these proprioceptive modalities individually has identified sense organ and afferent characteristics,^15,21,23–26^ premotor pathways processing proprioceptive information,^9,27–30^ and the role of individual modalities in motor control and locomotion.^7,10,16,31,32^

However, in most natural circumstances, load and movement occur simultaneously. During stance in walking, the legs both push against the ground (force) and move to propel the body. Simultaneous activation also occurs in still postures when external forces are large enough to move the originally stationary limb.^22,33,34^ Functionally correct responses thus require that premotor networks respond to movement in the context of load, and load in the context of movement, i.e., that the modalities be integrated in a context-specific manner.

Load and movement inputs converge in insect and vertebrate local networks^29,35^ and influence each other’s processing.^4,36,37^ In stick insects, simultaneous load alters movement reflexes^4,29^ and the occurrence of reflex reversals,^38^ in which resistance reflexes in standing animals become assistance reflexes during active movements.^39–41^ In stick insect legs, reflex reversal changes sensory processing at multiple levels of the motor network,^42,43^ suggesting a more complex integration of load and movement input than simple signal summation.

In stick insects, presynaptic inhibition between load (CS) and movement (femoral chordotonal organ (fCO)) afferents contributes to this interaction.^2^ However, these afferents also feed into a distributed network of premotor nonspiking interneurons (NSIs).^29^ How presynaptic inhibition affects network processing is unknown. We therefore traced the interplay of load and movement signals through the network.^27,29,33,42,44–46^ We hypothesized that simultaneous load would alter how the premotor network processed movement input, and result in movement reflexes being expressed in a load-context dependent manner. Because of the importance of gain explained above, although we describe the global changes in the input:output functions, we concentrate on changes in their gain.

Specifically, we investigated whether, in the femur-tibia (FTi) joint control loop, load input from tibial CS (tiCS) changed the gain of movement reflexes induced by fCO stimulation.^47^ This system is particularly well-suited for this investigation because tiCS synapse directly onto only one of the system’s three MNs, the fast extensor tibiae (FETi), but not the slow extensor tibiae (SETi) or common inhibitor 1 (CI1)^3^ and many of the premotor NSI’s onto which fCO and tiCS synapse are identified,^27,29,45^ as are the effects of these NSI’s on SETi activity,^27,47,48^ the MN which almost exclusively determines FTi joint angle during rest standing.^47^ We traced the interaction of load and movement signals from the earliest neuronal stage, presynaptic afferent inhibition, through the nonlinear summation and neuron-specific gain control in the premotor neural network, to the network’s output onto the extensor tibiae (ExtTi) MNs, and finally to the effects on muscle force. Motor output gain changes differed for different movement parameters and MN. Pharmacologically blocking presynaptic inhibition showed that it was essential for multimodal integration. By tracing multimodal signal interaction through the entire processing pathway—afferents, interneurons, and MNs—this work describes on the cellular and synaptic level how gain changes in a sensorimotor network can implement context-dependent processing of multimodal proprioceptive input.

## RESULTS

### Load (tiCS) and movement (fCO) sensorimotor pathways

tiCS and fCO afferents are major proprioceptive inputs in the FTi joint control system. In addition to fCO monosynaptic connections to MNs,^3,49,50^ tiCS and fCO afferents synapse onto premotor NSIs which in turn excite or inhibit SETi (Figure 1).^5,27,29^ Some NSIs also affect the fast ExtTi (FETi) or common inhibitor 1 (CI_1_) MN’s. When load or movement stimuli are presented individually, afferent firing, NSI membrane potential (V_m_), and MN firing accurately represent stimulus shape and parameters such as velocity or amplitude. Recordings from tiCS groups 6A and 6B (G6A, G6B), fCO, and downstream neurons during ramp-and-hold, sinusoidal, or triangle stimuli show the characteristic responses at each level of the pathway (Figure 1).

**Figure 1:**
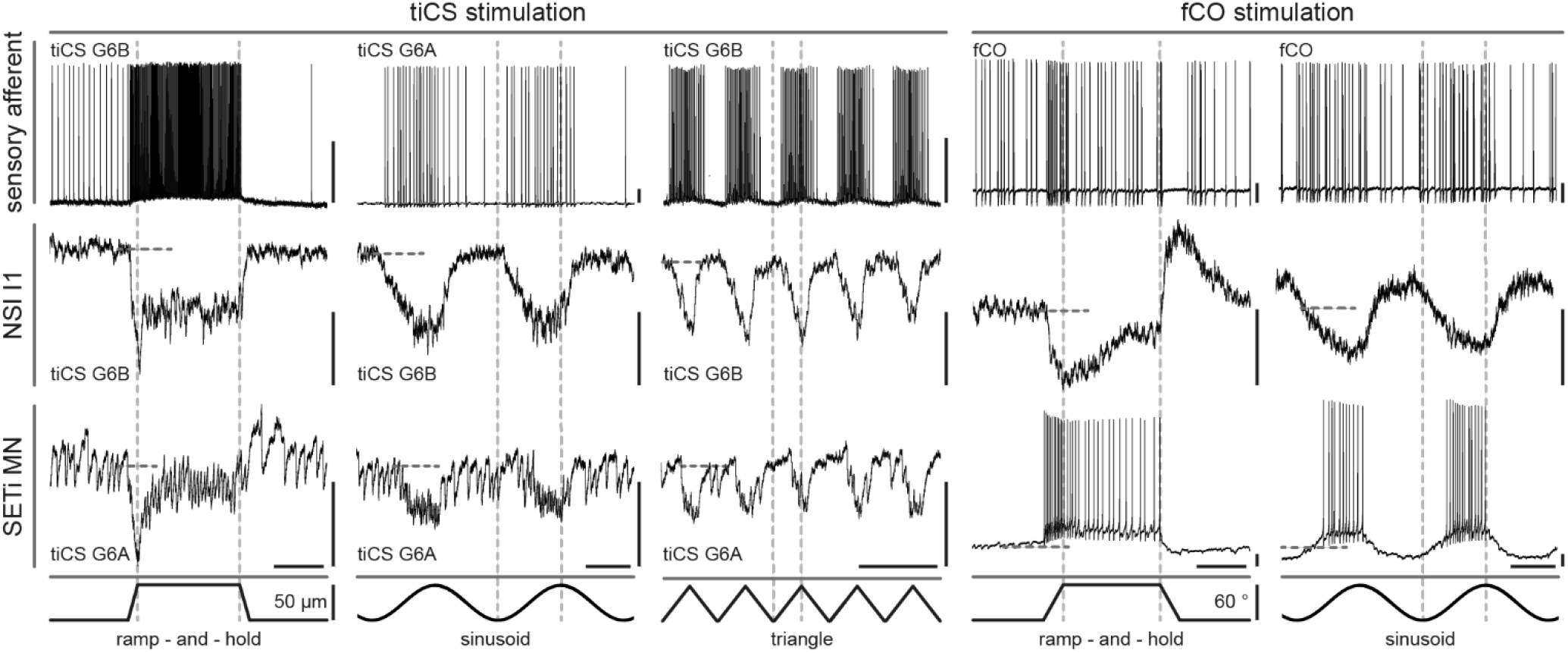
Intracellular responses to load (tiCS) and movement (fCO) applied individually. tiCS G6A and G6B (row 1, columns 1-3), fCO (row 1, columns 4, 5), an exemplary NSI (I1, 2^nd^ row), and SETi (3^rd^ row) responses to different mechanical stimulations (4^th^ row; upward deflections, increased G6A or G6B tibial bending (force) or fCO elongation (FTi joint flexion). Stimulated tiCS subgroup indicated in each panel in columns 1-3. Vertical dashed lines provided to facilitate comparing stimulation and responses. G6A and G6B responses had opposing directional sensitivity. Horizontal dashed lines: neuron resting V_m_. Scale bars: 0.5 s (horizontal), 5 mV (vertical).

### fCO afferents presynaptically inhibit each other and tiCS afferents presynaptically inhibit fCO afferents

Presynaptic inhibition between sensory afferents alters sensory input before it enters the premotor network.^1,10,51^ Our data confirm prior work showing that fCO afferents presynaptically inhibit one another.^52,53^ Recordings from fCO terminal neuropilar arborizations showed altered firing frequencies, primary afferent depolarizations (PADs), and reduced normalized action potential (AP) amplitude (0.985 ± 0.007, N = 5) during movement stimuli (Figure 2A). The latter two are hallmarks of presynaptic inhibition.^52,53^

**Figure 2.**
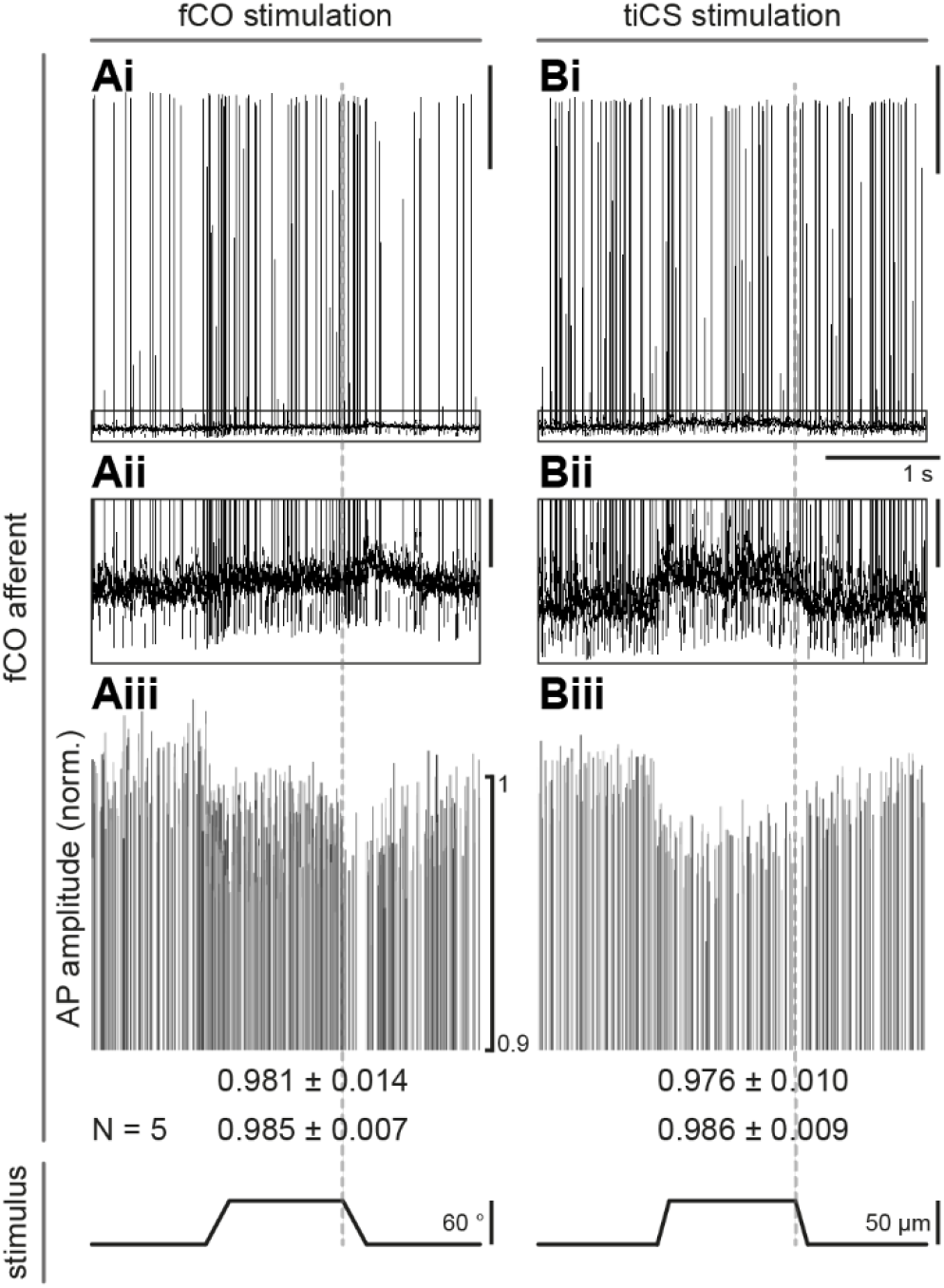
Presynaptic inhibition between fCO afferents and of fCO afferents by tiCS afferents. Intracellular recording of an fCO afferent during fCO (A) and tiCS (B) stimulation. fCO AP frequency was altered by fCO but not tiCS stimulation (i). Both stimuli elicited PADs (ii, enlarged view of inset shown in i) and reduced AP amplitude (iii). Data in iii normalized to average AP amplitude in 1 s interval before stimulation. All data from same fCO afferent, 5 consecutive stimuli overlaid in each panel. Bottom numbers: reduction of AP amplitude for this neuron (top) and for N = 5 (bottom, pooled results of G6A and G6B stimulations). Scale bars: 20 mV (i), 2 mV (ii). Aiii scale bar applies to Biii, Bi time scale applies to all panels.

Load and movement sense organs are rarely activated separately in natural conditions. tiCS stimulation in the same preparation as in Figure 2A did not alter fCO firing frequency but did elicit PADs and reduce normalized fCO AP amplitude (0.986 ± 0.009, N = 5, Figure 2B). tiCS effects on fCO afferents were thus due to presynaptic afferent inhibition. Movement and load-elicited PADs outlasted the stimulus (vertical dashed lines), and effects on AP amplitude could outlast the PADs, as has been observed in other preparations.^1,54^

### fCO and tiCS inputs non-linearly summate in identified premotor network neurons

fCO and tiCS afferents mono- and polysynaptically connect onto individually identifiable premotor NSIs.^3,27,29,45^ tiCS presynaptic inhibition of fCO responses would be expected to alter NSI processing of fCO and tiCS input when both modalities are active. We therefore measured NSI responses to sinusoidal stimulation of fCO and tiCS alone or together (Figure 3A top row). The combined responses (blue traces) differed from algebraic summation (grey traces) of the individual responses, indicating that the summation was nonlinear (Figure 3A bottom row, total N = 20). This non-linearity is particularly well shown by NSI E2 (Figure 3Aii), in which load stimuli alone did not elicit a response, but activation of both load and movement resulted in a response different from that elicited by movement alone.

**Figure 3:**
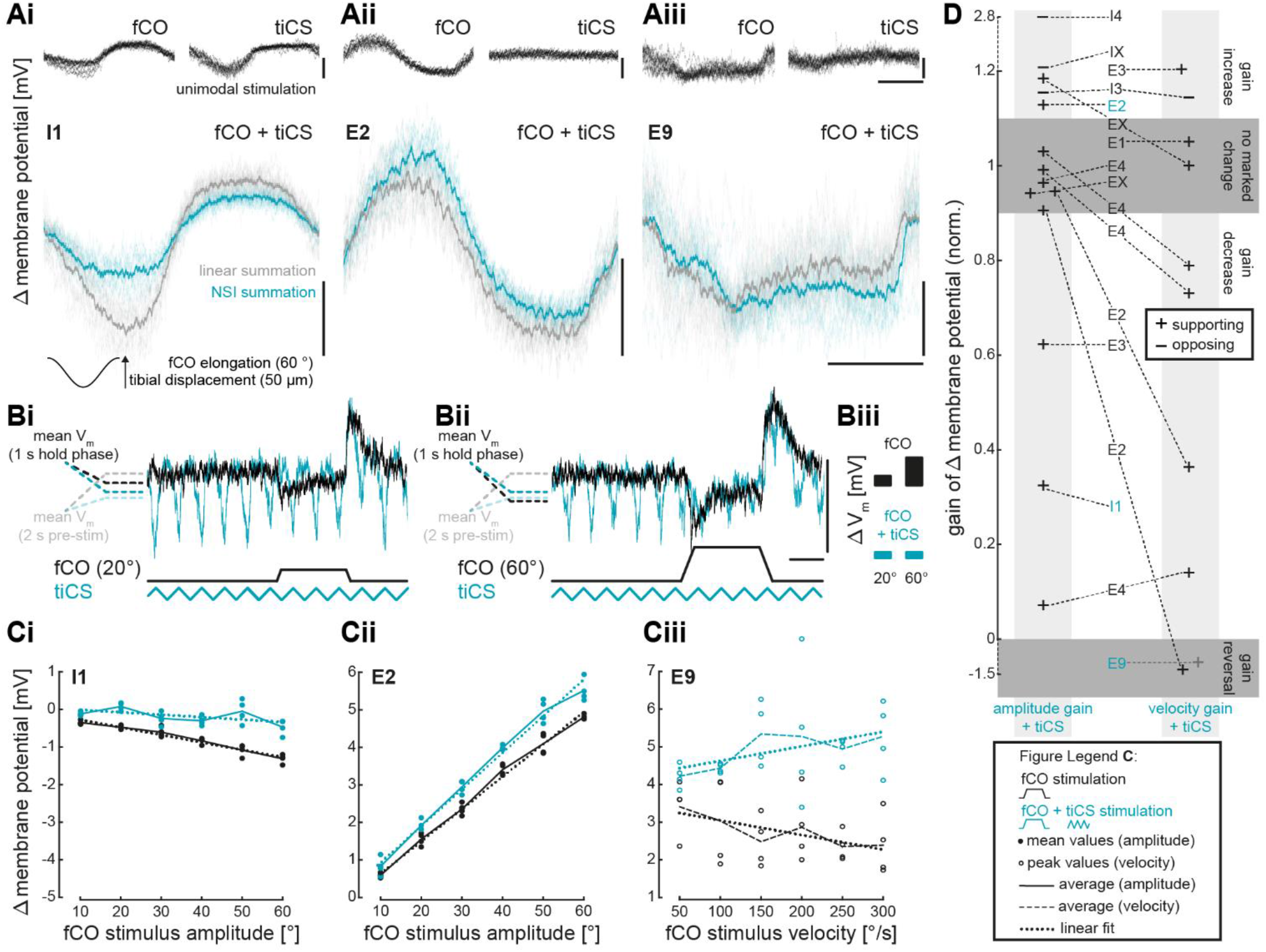
Nonlinear summation and neuron-specific gain modulation in NSIs. A: Load (tiCS) and movement (fCO) signals summed nonlinearly in NSIs. Data from NSIs I1 (Ai), E2 (Aii), and E9 (Aiii). Top row (black): NSI V_m_ response to sinusoidal fCO (left) or tiCS (right) stimulation alone; 15 overlaid consecutive stimuli shown per panel. Bottom row: light blue, NSI response to combined fCO and tiCS sinusoidal stimulation; grey, algebraic sum of responses to individual stimuli in top row (15 overlaid stimuli in both cases). Dark lines in bottom row: mean traces. Vertical scale bars, 5 mV; horizontal, 0.5 s. B, C: tiCS stimulation (blue triangle wave, B) altered NSI fCO response gain. B: NSI I1 response to ramp and hold fCO stimuli without (black) and with simultaneous tiCS activity (blue) at two fCO hold amplitudes, scale bars as in A. Panel Biii quantitatively compares responses to fCO stimulation at two amplitudes without (top row) and with (bottom row) tiCS stimulus. See text for detailed explanation. C: NSI gain (slope of linear fits) dependence on hold amplitude (i, ii) and velocity (iii) (black). Gain was decreased (i), increased (ii) or reversed (iii) by tiCS stimulation (blue). Inset: key to point and line identifications. D: NSI classification as functionally supporting or opposing fCO movement reflex (+, −) and gain change to the amplitude (left column) or velocity (right column) components of the fCO ramp-and-hold stimulus caused by tiCS stimulation. E, I in center column indicate excitatory or inhibitory effects on SETi; X indicates the NSI could not be identified by type (2, 3, 4, etc.). Blue names in central column identify neurons shown in A-C. Data excluded if R^2^ of linear fit < |0.6|. See STAR Methods for how gain was calculated. N (number of NSIs) in D = 17.

### Neuron-specific gain modulation in the premotor network

The non-linear summation shown in Figure 3A suggested that, in addition to changes in overall response amplitude (e.g., the global depolarization of NSI E2 in Figure Aii), which depend on the activated tiCS subgroup (G6A, G6B),^29,55^ that tiCS input could also be altering the gain (the ratio between signal input and output) of NSI responses to fCO input. We quantified these gains using ramp-and-hold stimuli as follows. Figures 3Bi and 3Bii show that NSI I1 response (hyperpolarization) to fCO ramp-and-hold stimuli increased with hold amplitude. The gray and black dashed lines in 3Bi and 3Bii show, respectively, mean I1 V_m_ in the 2 s prior to fCO stimulation and during the stimulus. The distance between the gray and black lines is clearly larger for the larger hold amplitude (panel Bii); this response difference is shown by the black bars in panel Biii. NSI responses to changes in hold amplitude (and on-ramp velocity), in the ranges applied here, are approximately linear.^27,45,56^ Linear fits (dashed lines in Figure 3C) were therefore applied to all NSI response-amplitude and response-velocity data. NSI gain is the slope of these fits (for I1, Figure 3Ci, black trace).

The blue traces in Figure 3Bi and 3Bii show NSI I1 responses to fCO ramp-and-hold stimuli with a simultaneous tiCS triangle stimulus. tiCS input caused I1 hyperpolarizations phase-locked with each peak of the triangle stimulus. I1 V_m_ before fCO stimulation was therefore more hyperpolarized than in the absence of tiCS stimulation (difference between gray and light blue dashed lines, 3Bi and 3Bii). If the two inputs summed algebraically, the result in 3Bi would be a global hyperpolarizing offset of the tiCS rhythmic hyperpolarizations equal to the (black) fCO response without tiCS stimulation. No such translocation occurred. fCO stimulus therefore changed mean I1 V_m_ only slightly (light and dark blue lines in 3Bi, blue bars in 3Biii). For the large fCO stimulus (3Bii), I1 response to tiCS stimulation was complicated, with the initially most obvious change being a decreased tiCS oscillation amplitude, likely because fCO-induced hyperpolarization shifted V_m_ closer to the reversal potential of tiCS-elicited hyperpolarizations. However, determining how tiCS stimulation affects the gain of I1 to fCO input requires comparing mean I1 changes with and without tiCS stimulation. In 3Bii, tiCS stimulation alone again considerably hyperpolarized mean I1 V_m_ (light blue line). fCO stimulus only slightly changed this mean hyperpolarization (compare dark and light blue lines in 3Bii and right column of 3Biii). tiCS stimulus thus decreased I1 response to fCO input at this particular fCO input value. Demonstrating that tiCS changed I1 response gain, however, requires plotting fCO responses without and with tiCS input across a range of fCO input values, since the gain of the fCO to I1 synapse is the slope of this input:output function (Figure 3Ci). The blue fit in Figure 3Ci shows that tiCS indeed decreased I1 gain (slope) to fCO input. In I1, the change was almost completely a gain change, as I1 responses to low fCO stimulus amplitudes changed only slightly.

Figure 3Cii shows that NSI E2’s response (depolarization) also increased with hold amplitude. However, in E2 simultaneous tiCS input increased the gain of E2’s response to fCO input. Again, tiCS’ effect was almost exclusively a gain change, as tiCS stimulation again only slightly changed I1 response at low fCO values. Figure 3Ciii shows NSI E9’s response to fCO input alone was a decreasing depolarization, in this case to the slope (velocity) of the ramp and hold stimulus (see STAR methods for how response-velocity data were quantified). For E9, tiCS stimulation induced both effects described in the Introduction. First, it globally translocated the data to more positive values (at all fCO values, E9 depolarization was increased). Second, in addition to these frank changes in synaptic strength, it again affected synapse gain, reversed it from negative to positive. The data translocation will affect network activity, as is more fully described below (see Figure 4). However, because it does not affect the sensitivity of the network to input, this effect has less effect on network stability than does the gain change. We therefore here concentrate on tiCS-induced gain changes. Figure 3C thus shows that tiCS modulation of NSI gain to fCO stimuli depended both on NSI identity (e.g. I1, E2, E9) and movement parameter (amplitude, velocity).

**Figure 4:**
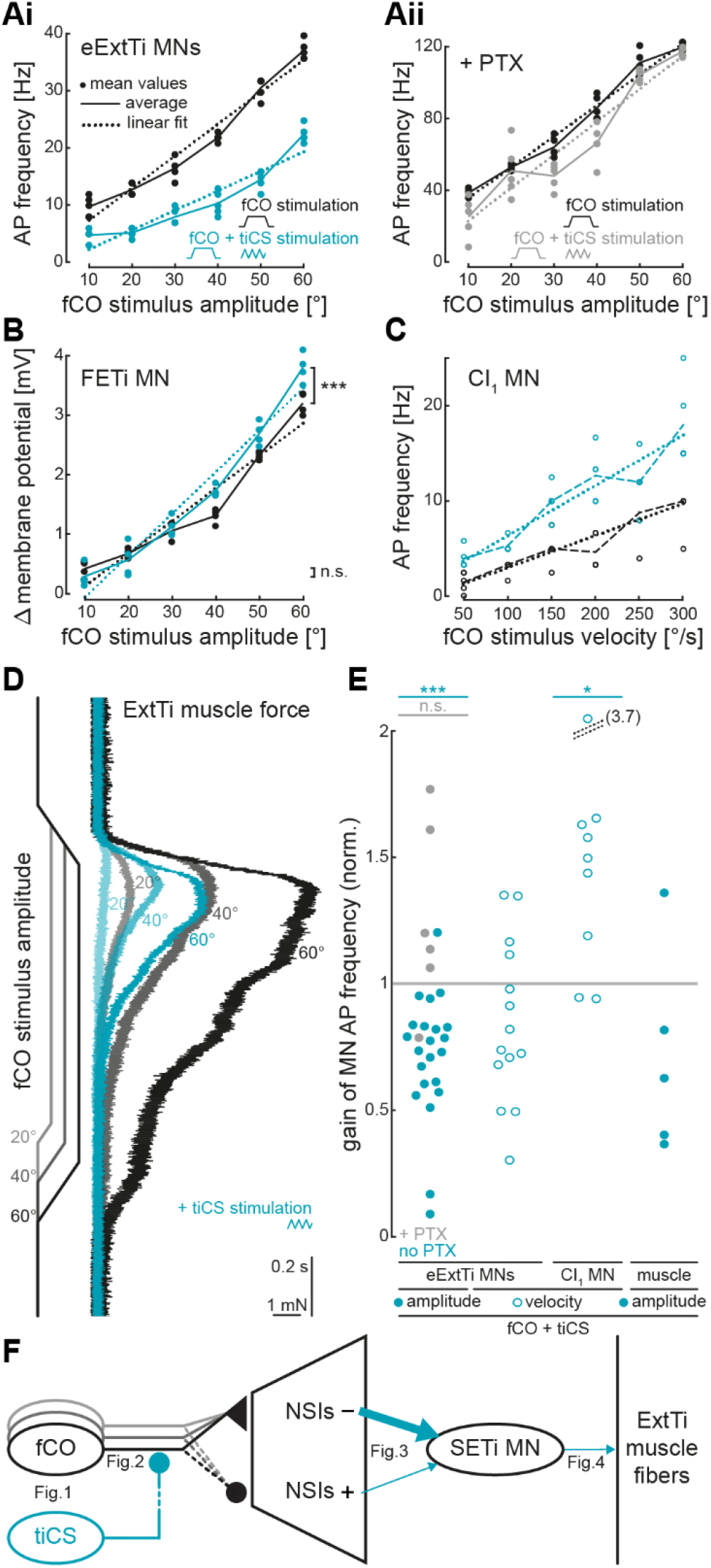
Gain modulation of ExtTi MN activity and ExtTi muscle force. A: tiCS stimulation reduced eExtTi MN overall activity and gain (slopes of blue and black linear fits) to the fCO amplitude component, measured using AP frequency. Both effects were abolished when presynaptic inhibition was blocked with picrotoxin (PTX, Aii). B: FETi V_m_ significantly increased at high (60°), but not low (10°) movement stimulus amplitudes, resulting in a gain increase. C: CI_1_ overall activity and velocity-dependent gain, measured as AP frequency, increased. A, B, C: Data from single animals, key same as inset in Figure 3. D: tiCS reduced amplitude of isometric ExtTi muscle force (compare black and blue contractions) and decreased gain. E: Grouped data. 1^st^ and 2^nd^ columns: tiCS-induced changes of eExtiTi MN gains to fCO amplitude and velocity components. 3^rd^ column: tiCS-induced change in CI_1_ gain to fCO velocity component. 4^th^ column: tiCS induced change in ExtTi gain to fCO amplitude component. All data normalized to exclusive movement stimuli. Filled and open circles, data from single animals, amplitude and velocity, respectively. Grey: PTX trials. F: Schematic of how tiCS presynaptic inhibition alters movement amplitude gain. Multiple fCO inputs symbolize how different effects on fCO and tiCS inputs on individual NSI’s results in net increase in fCO reflex-opposing NSI network output. Dashed lines: polysynaptic pathways; triangle, effective excitatory input; circle: effective inhibitory input; +/-, fCO reflex supporting or opposing NSI, respectively.

Depolarization of individual NSI increases or decreases SETi activity (hence their identification as E or I neurons).^27,45^ Combining these identifications with NSI response to fCO input (depolarization, hyperpolarization) allows classifying each NSI as supporting or opposing the movement reflex (“+”“-” signs in Figure 3D).^27,45^ For example, I1 depolarization inhibits SETi. However, I1 hyperpolarizes in response to the ramp-and-hold stimulus, thus functionally supporting the movement reflex. Figure 3D summarizes for the NSI’s examined here (names in central column) whether tiCS stimulation increased or decreased the neuron’s gain, sorted by which component (amplitude, left column; on-ramp velocity, right column) of the ramp-and-hold fCO stimulus was being examined. For NSIs for which an entry is present in only one column, the NSI could not be recorded from long enough to measure responses to both components.

The vertical position of each NSI’s symbol (+, −) shows the gain change caused by tiCS stimulation. Thus, tiCS stimulation strongly decreased I1 gain to the hold component of the fCO stimulus (the “+” in the left column connected by the dashed line to the “I1”), to 0.32 normalized to its fCO-exclusive gain (1 indicates no change in gain). The dark rectangle at the bottom of the panel highlights the cases, as in Ciii, in which tiCS input reversed NSI gain. In none of these cases, however, did this gain reversal change the neuron’s effect on SETi (e.g., in 4Ciii, in which tiCS simulation reversed the gain, the effect on SETi was still supporting at all fCO stimulation amplitudes).

Examining these summary data showed three properties. First, responses of neurons of the same NSI type could vary. For instance, tiCS stimulation did not affect gain for three E4’s (those in left column of upper gray horizontal rectangle) but strongly decreased the gain of a fourth (left column “+” E4 near bottom of Figure 3D). Second, for the large majority (9 in the amplitude column, 6 in the velocity column, out of a total of 20 “+” measurements in both columns), tiCS stimulation either did not affect or decreased the gain of NSI’s supporting the movement reflex. Third, tiCS stimulation increased the gain of all NSI’s opposing the movement reflex (“-” entries). The net effect of the NSI network on SETI caused by tiCS activity was thus a decrease in inputs supporting the movement reflex and an increase in inputs opposing it.

### Parameter-dependent and neuron-specific modulation of tibial MN gain

Three MNs, SETi, FETi, and CI_1_, innervate the ExtTi muscle. In the resting animal, SETi mediates posture control. FETi is recruited during active movements.^47,48,57^ CI_1_ activity also increases during active movements, presumably to decrease the contraction amplitude and increase the relaxation rate of the slowly-relaxing SETi component of the ExtTi contractions. In posture, SETi activity constitutes the majority of excitatory ExtTi (eExtTi) MN APs during a movement reflex response, with FETi AP numbers, especially at lower stimulus amplitudes and velocities, lying in the low single-digit range.

The NSIs affect SETi, FETi, and CI_1_ activity differently. It was therefore important to test tiCS effect on each MN (Figure 4). In these experiments we used the same ramp-and-hold fCO stimulation paradigm as in Figure 3. As with E9, tiCS stimulation changed both the overall response (global translocation of blue data points to lower AP frequency in Figure 4Ai), which depended on the stimulated subgroup of tiCS^55^, and the gain (slope decrease) of eExtTi MN (almost exclusively SETi) responses to fCO stimuli, which was independent of tiCS subgroups (data from one experiment shown in Figure 4Ai; all experiment data in Figure 4D, 1^st^ column, blue, N = 22, p = 4.8 x 10^-6^). Both effects depended on presynaptic inhibition, as they were abolished by picrotoxin application (Figure 4Aii, N = 1, Figure 4D, 1^st^ column, grey, N = 6, p = 0.14), a GABA_A_ antagonist that specifically blocks presynaptic inhibition in the stick insect premotor network.^52^ tiCS simulation did not consistently alter gains of the eExtTi MN velocity-dependent responses (individual experiment data not shown, all experiment data shown in Figure 4D, 2^nd^ column, N = 14, p = 0.09).

Because of the very low FETi firing rate in the inactive animal, we measured the changes in FETi V_m_ to fCO and tiCS stimulations (Figure 4B), and in only two experiments. Consistent with FETi playing a negligible role in the movement reflex during posture, FETi’s response to hold amplitude was very small, reaching a maximum of 3 mV at the largest hold amplitude. At this amplitude, simultaneous tiCS stimulation significantly, but slightly (1 mV), increased FETi depolarization. tiCS also increased FETi gain (compare slopes of blue and black lines in Fig. 4B; p = 7.5 x 10^-4^ for the data from both experiments). tiCS stimulation had no effect on FETi response to the velocity component of the ramp and hold fCO stimuli.

The inhibitory ExtTi MN, CI_1_, does not respond to the hold component of the fCO stimulation.^47,48^ It does respond to fCO velocity, increasing with increased velocity (Figure 4C, black). Simultaneous tiCS input again affected both CI_1_ overall response (upward translocation of all data points) and gain to fCO velocity (individual experiment data, Figure 4C; all experiment data, Figure 4D, 3^rd^ column, N = 9, p = 1.1 x 10^-2^).

Understanding the effect of tiCS stimulation on ExtTi muscle response to fCO stimulations requires considering both the effects on overall activity and gain. tiCS simulation reduced ExtTi contraction amplitude at all tested fCO amplitudes (Figure 4D, black traces).^58^ These amplitude changes are almost certainly due to the overall changes in eExtTi (primarily SETi, Figure 4Ai) responses caused by simultaneous tiCS activity (because velocity was not altered in these experiments, changes in CI_1_ activity did not play a role), not changes in eExtTi gain. However, the changes in eExtTi gain would nonetheless be expected to change the gain of ExtTi muscle responses. We therefore calculated ExtiTi gain in the same way as for NSIs and MNs, by measuring the contraction amplitudes, plotting them against fCO stimulation angle, and applying linear fits to the data. Although the gain changes did not achieve statistical significance (p = 0.42), Figure 4E, 4^th^ column shows that in four out of five experiments, muscle gain to fCO input indeed decreased.

Figure 4F summarized our results, showing how tiCS presynaptic inhibition of fCO afferents, as non-linearly processed by the NSI network, result in a net gain increase in input to SETi of movement reflex opposing NSIs, and thus a decrease in ExtTi muscle amplitude and gain.

## Discussion

Understanding motor systems requires understanding how simultaneous inputs from different sensory modalities are processed by sensorimotor networks. In many systems, inability to trace activity through neural pathways on the cellular level limits our ability to achieve this goal. In the stick insect, individual neurons of the FTi joint motor control neural network can be recorded from, and sensory input precisely activated with mechanical stimuli imicking those that occur naturally. We followed here interactions between proprioceptive load (tiCS) and movement (fCO) sensory input from afferent to muscle output levels. fCO inputs presynaptically inhibited each other and were presynaptically inhibited by tiCS afferents. Both types of afferents feed, in part monosynaptically, into the FTi premotor NSI network. In the presence of load, nonlinear signal summation and differential sensory gain modulation within the network altered the overall responses and gains to movement stimuli. With respect to the gain alterations, load decreased the gain of NSI neurons supporting, and increased the gain of NSI neurons opposing, the movement reflex response. SETi and isometric ExtTi gain to movement sensory input consequently decreased. These results demonstrate how load input tunes movement reflex pathways and provides a cellular basis for context (load) dependent expression of the movement reflex.

### Limitations in interpretation of these data

Although the gross effects of the NSI’s on SETi activity have been described, the NSI’s being non-spiking means that monosynaptic connections in the NSI network, and to SETi, have not. However, because the overall effect of each NSI on SETi is known, this lack does not undermine the conclusions drawn here. NSI E4 showed a range of changes in response gain. This is likely because there are at least two types of E4 NSI’s.^59,60^ Because we analyzed the data from all neurons recorded from individually, this result again does not undermine the conclusions drawn here.

#### Comparison with prior work

Follmann et al. examined integration of mechano- and chemosensory sensory input in the crustacean stomatogastric system using optical techniques to determine which neurons were activated by the two modalities alone and together in the commissural ganglion to which these afferents project.^61^ This work showed that the multimodal information was represented by a non-linear combinatorial code, in that the neurons active when both modalities were stimulated differed from the sum of those activated by each modality alone. This differs from the stick insect NSI network, in which the activity of almost all neurons is altered by load or movement stimulation applied alone.^27,29^

The use in Follman et al. of optical methods (as opposed to intracellular recordings), the fact that the commissural ganglion neurons are not identified, and the lack of stimuli with incremental parameter changes (e.g., increasing amplitudes) precludes identifying presynaptic inhibition or other inter-afferent interactions, and analyzing gain changes in commissural ganglion pathways. The data presented here thus represents a qualitative advance in understanding how multimodal integration occurs.

### Proprioceptive integration in premotor NSIs

The stick insect premotor NSI network processes movement and load input in a distributed, antagonistic manner, i.e., multiple parallel pathways that support or oppose the motor output being generated^27,29,44,45^ In the context of reflex reversal and active behavior, shifting the weighting of these pathways as a mechanism to control MN gain has been suggested, but never directly shown.^42,47,62–65^ We present here such evidence. Controlling sensory input gain in distributed networks is necessary to use the same network for different tasks and to support behavioral flexibility. Distributed processing is a central component of the mechanism presented here; it allows the system to fine-tune individual network pathways and modulate expression of one sensory modality as a function of another using a single set of NSI pathways.

### Functional relevance of sensorimotor gain control

Presynaptic inhibition is ubiquitously (locust^14^ to monkey^66^) used to control sensorimotor gain.^1,51,67^ It is essential for stable motor output,^10^ to avoid sensory habituation and saturation,^53,68^ and to ensure proper coordination during walking.^13^ By controlling the gain of sensory signals, the sensitivity to these inputs of downstream neural networks is up- or downregulated. Modulation depends on signal type, relevance for behavioral context, and need to stabilize the system and prevent signal overload.^11^ Our detailed analysis of differential fine-tuning of individual interneuronal and MN pathways show how a small locomotor network can implement context-dependent sensorimotor control. FTi joint motor control system movement signal processing and gain differed in the presence or absence of load. Because changes in load and movement frequently occur simultaneously, these results underline the necessity of considering proprioception as a multimodal sensory input.

The stick insect ExtTi muscle being innervated by only three MNs, the fast and the slow excitatory FETi and SETi and the inhibitory CI_1_,^47^ facilitates functional analysis of the proprioceptive integration demonstrated here. In the standing animal, SETi firing helps set FTi joint angle. The fCO reflex pathway, when activated alone, increases SETi firing with joint flexion, and thus is a classic resistance reflex that acts to maintain a constant joint angle. When load is simultaneously applied, the gain of SETi activation by joint flexion is reduced. This seems self-defeating, as it decreases the ability of the fCO reflex to maintain FTi joint angle. However, at some load magnitude, maintaining a specific angle would require forces that would damage the FTi joint complex. Thus, fCO and tiCS would work together to 1) at relatively low levels of load (low or no tiCS activation) allow fCO to maintain joint posture and 2) at dangerous load levels (high tiCS activation) reduce the fCO reflex to allow joint rotation. Reducing gain is a particularly appropriate method to achieve this goal because its reductions become larger with increased load, as opposed to a simple reduction (translocation) of the input:output curve. Reducing MN gain to appropriate sensory inputs could thus be a general mechanism to prevent damage as resisted external forces become too large. Consistent with this interpretation, tiCS are believed to tune FTi muscle activities when their shortening when activated is resisted (which activates tiCS).^69^

FETi and CI_1_ are recruited during active movements, with CI_1_ presumably active in part to decrease ExtTi slow fiber contraction and increase slow fiber relaxation. In the work presented here, SETi velocity-dependent gain was not consistently altered, whereas CI_1_ velocity-dependent gain increased, and FETi amplitude-dependent gain showed a tendency towards increase. These changes are consistent with the decreased SETi activity and increased CI_1_ and FETi activity that occur when animals switch from standing to active movements^70^, i.e., to a reversal of the fCO reflex, and with data showing that load input supports this reversal.^39^ They are also consistent with the switch to the active state beginning with decreased SETi and increased CI_1_ activity.^39,71^ Interestingly, in a model based on the stick insect premotor network, reflex reversal could be evoked by altering the gain of specific movement afferent pathways,^72^ a mechanism not identical, but surprisingly close, to the one presented here.

Until recently, load and movement input were believed to be processed primarily by separate pathways. We recently demonstrated that, in fact, load and movement signals are processed within the same NSI network,^29^ albeit with different temporal characteristics.^3^ Load processing is delayed relative to movement, resulting in load being optimally timed to tune ongoing movement processing. Here, we examined simultaneous load and movement activation and showed that load altered a movement reflex as a function of (in the context of) load. In a study on the convergence of touch and proprioceptive signals in the ventral nerve cord of *Drosophila melanogaster*, Tuthill and Wilson discuss putative contextualization of load and movement input, which we have demonstrated here.^37^ Future studies will show whether the contextualization of sensory signals by means of sensorimotor gain control is a general mechanism of motor networks across animals.

## Acknowledgements

The authors thank Till Bockemühl for helpful discussions on data analysis and Mehrdad Ghanbari and Michael Dübbert for technical support. This study was supported by a grant from the Deutsche Forschungsgemeinschaft (DFG, German Research Foundation) 233886668/ GRK1960 to AB and CG and supported by the Studienstiftung des deutschen Volkes to CG.

## Author contributions

CG and AB conceived and designed research; CG performed experiments, analyzed data, and prepared figures; CG,+ AB, and SLH interpreted results; CG drafted manuscript; CG,AB, and SLH edited and revised the manuscript and approved the final version.

## STAR Methods

### Lead Contact

Further information and requests for study-related resources should be directed to the lead contact, Corinna Gebehart.

### Materials Availability

No unique reagents were generated in this study.

### Data and Code Availability

Source data and code used for analysis are available upon request to the lead contact, Corinna Gebehart.

### Animals

Experiments were conducted on adult female stick insects, *Carausius morosus*, from the parthenogenetic colony at the University of Cologne. The colony is kept at 22 – 24 °C, 50 % humidity, 12 h light / dark cycle, and fed with blackberry leaves.

### Dissection

Animals were dissected as described in ^29^. In short, all legs except for the right mesothoracic leg were removed; the thorax-coxa, coxa-trochanter, and FTi joints were immobilized at 90 ° angles using light curing glue (3M ESPE Sinfony, Neuss, Germany) and a minuten pin inserted into the ventral trochanter. The animal was pinned to a platform, the distal femur glued to the edge with the tibia protruding from the platform. The mesothoracic ganglion was exposed by a dorsal midline incision; all lateral nerves except for the ipsilateral nervus cruris and lateralis 3 were squeezed (nomenclature according to ^73^).

For intracellular recordings, the ganglion was pinned with cactus spines on a wax-coated steel platform, cf. ^27,74^. The ganglion sheath was softened with Pronase E crystals (VWR, Darmstadt, Germany). The body cavity was rinsed and filled with extracellular saline (178.54 mM NaCl, 17.61 mM KCl, 7.51 mM CaCl_2_, 25 mM MgCl_2_, 10 mM HEPES buffer, set to pH 7.2 with NaOH).^75^ All experiments were performed in inactive, resting animals.^47^

### Electrophysiological recordings

Intracellular sharp microelectrodes (35-45 MΩ) were fashioned from borosilicate glass capillaries (8GB100TF-8P, Science Products, Hofheim, Germany) using a micropipette puller (P-1000, Sutter Instruments, Novato, CA, USA) and contained chlorinated silver wire. Electrodes were filled with intracellular saline (1M KAc, 0.1 M KCl). 5 % neurobiotin tracer (Vector Laboratories Cat# SP-1120-20, RRID:AB_2336606) was added for morphological neuron identification, for staining and fixation procedure, see.^29,76^

Intracellular signals were recorded from arborizations of NSIs and MNs in the medio-dorsal neuropil of the ipsilateral hemiganglion, and from afferent arborizations at the point where nervus cruris enters the ganglion. Intracellular signals were compensated for interstitial potential (5-10 mV),^77^ and amplified (gain x20, highcut filter 3 kHz, SEC-06 preamplifier, SEC-10LX single electrode clamp amplifier, npi, Tamm, Germany). Extracellular signals of ExtTi MNs were recorded with a hook electrode from nerve F2 within the femur,^78^ and amplified (gain x2000, lowcut filter 0.3 kHz, highcut 2.5 kHz, notch filter 50 Hz, MA101 preamplifier & MA102 differential amplifier, Electronics Workshop, University of Cologne). All signals were digitized and stored with Spike2 software (sampling rate 12.5 kHz for extracellular, 6.25 kHz for intracellular signals, ADC Micro1401 mkI, Spike2 v7.16, RRID:SCR_000903, CED, Cambridge, UK).

### Mechanical sensory stimulation

The fCO was stimulated by opening the dorsal proximal femur, clamping, and moving the receptor apodeme.^79,80^ Receptor elongation mimics tibial flexion (upward deflection of stimulus trace in the following figures), vice versa for receptor elongation. Ramp-and-hold fCO stimuli in experiments where stimulation amplitude was altered had a consistent ramp velocity of 300 °/s. Ramp-and-hold fCO stimuli where stimulation velocity was altered had a consistent hold amplitude of 60 °, corresponding to 300 μm elongation of the receptor apodeme.^81^

tiCS were stimulated by a metal probe that pushed the proximal tibia into a ventral or dorsal direction by 50 μm in all stimulus paradigms, activating groups 6A or 6B (G6A, G6B), respectively.^82^ In the following figures, an upward deflection of the stimulus trace denotes an increase in load, a downward deflection the return to control condition without load stimulus. Ramp-and-hold tiCS stimuli had rise and fall times of 0.1 s each, with a hold phase of 1 s in between. Triangle tiCS stimulus rise or fall ramp duration was 0.17 s without a hold component to avoid adaptation and was used as a continuous stimulation in experiments with fCO stimuli of increasing amplitudes or velocities. Control intracellular recordings of tiCS afferents did not show adaptation to triangle stimuli of the duration applied in this study (N = 2). The order of exclusive movement and combined load and movement stimuli was varied to exclude adaptational effects. We controlled for habituation to the ongoing tiCS triangle stimuli by presenting the stimulus paradigm to load-habituated animals. Habituation did not affect the reported changes in gain.^83^

Sinusoidal waveform stimuli of the fCO and tiCS had a frequency of 0.7 Hz, a maximum amplitude of 60 ° or 50 μm, respectively, and consisted of 15 continuous cycles. Intervals between all types of stimuli were at least 2.6 s long. The time within stimulation paradigms between different stimulus velocities or amplitudes was at least 10 s during which neither tiCS nor the fCO were stimulated. Mechanical stimulators were controlled by Spike2 or a digital stimulator (MS501) and moved by custom-built linear motors (lowpass filter 1 kHz, VCM Controller/Power Amplifier, Electronics Workshop, University of Cologne).

### Muscle force measurement

ExtTi isometric muscle contraction force was measured with a force transducer (ALS, Aurora 300 B dual-mode lever system; Aurora Scientific Inc., Ontario, Canada, resolution 0.3 mN), digitized, and stored (sampling rate 6.25 kHz, ADC Micro1401 mkl, Spike2, CED). ALS force level was set above the maximum muscle force to maintain constant muscle length. The most distal, dorsal part of the femur was cut open. The cuticular attachment site of the ExtTi muscle tendon was detached from its insertion in the FTi joint and fixated with a hook-shaped minuten pin that was attached to the ALS lever arm. Force offset was determined with the pin in position before the muscle tendon was attached. Stimulus runs with FETi MN activity were excluded from the analysis.

### Pharmacology

To block presynaptic inhibition, the ganglion was superfused with the noncompetitive GABA_A_ (γ-aminobutyric acid A) receptor antagonist picrotoxin (Sigma-Aldrich).^52^ Starting concentration was 0.03 mM, incomplete removal of extracellular saline from the body cavity prior to drug application led to further dilution. An increase in ExtTi MN frequency was used as an indicator for the effect of picrotoxin. Stimulations were started when MN frequency had reached a constant level, approx. 10 min following drug application. Both stimulus paradigms, i.e. movement stimuli with or without simultaneous load stimulation, were analyzed before and after application of picrotoxin and compared within each condition. Picrotoxin effects on the gain of velocity dependence were not tested; picrotoxin abolishes velocity dependence in the system, potentially because blocking presynaptic inhibition causes the gain of velocity information in the system to saturate.^52,84^

### Neuron identification

Local premotor NSIs were identified as nonspiking if no APs were elicited by current injection, tactile or sensory stimulation, and by subthreshold effects on MNs.^85^ Individual types of NSIs were identified based on their excitatory or inhibitory effects on ExtTi MNs, and their responses to fCO ramp-and-hold stimuli.^27,86^ Afferents were identified by their responses to sensory stimuli. Current injection into the neuropilar arborizations of these neurons fails to alter their AP frequency. ExtTi MNs were identified by the time-locked relationship between intracellularly and extracellularly recorded AP’s. Identities of the different types of neurons were verified by their morphology.

### Data Analysis

Activation of tiCS subgroups G6A and B by ventral and dorsal tibial bending, respectively, has differential effects on the firing rate of ExtTi MNs, i.e. inhibition by G6A and excitation by G6B.^55^ The subgroups showed no differential effects on the gain of NSIs and ExtTi MNs, therefore results were pooled. For linear summation of NSI responses to sinusoidal stimuli, the offset of 15 sweeps of responses to exclusive fCO and tiCS stimulation was removed and response traces to fCO and tiCS stimuli were summed.

Changes in gain of neuron activity or muscle force were tested by incremental increases of fCO ramp-and-hold stimulus amplitude or velocity with or without concurrent triangle stimuli to tiCS.

The initial ramp-and-hold fCO stimulus of each tested amplitude or velocity started simultaneously with the triangle tiCS stimulus, which, in experiments with intracellularly measured changes in V_m_, consistently led to outliers and was thus discarded. Only responses to the following 4 ramp-and-hold stimuli were included in these analyses. This effect was not observed in experiments where extracellularly measured AP frequencies were analyzed, thus all 5 ramp-and-hold stimuli were included.

Changes in V_m_ to increasing stimulus amplitudes were measured as the average change from resting potential during the hold phase of the fCO stimulus. For increasing stimulus velocities, depending on the overall response direction of the neuron, the maximum or minimum change from resting potential during the rising ramp of the fCO stimulus was analyzed. For analysis of AP frequencies, we used the average frequency during the hold phase (increasing stimulus amplitude) or the rising ramp (increasing stimulus velocity). We did not distinguish between AP’s from the slow and fast ExtTi MNs (SETi, FETi) in analysis of extracellular activity, since suprathreshold responses of FETi to fCO ramp-and-hold stimuli were much sparser than those of SETi, thus the bulk of the effect will be based on SETi activity, with FETi having little to no effect on the analyses. Within one recording, only stimulus runs with similar baseline levels of MN firing frequencies were compared. APs of the common inhibitor 1 MN (CI1) were extracted from extracellular F2 recordings by spike sorting. CI_1_ is not reliably active during the hold phase of ramp-and-hold stimuli, thus only its velocity-dependence during ramps could be analyzed.

We chose linear fits for our data, as they yield a more intuitive read-out of the slope, i.e. gain, of a curve than sigmoidal curves, although the latter typically is more realistic for most physiological data. The parameters tested were chosen because they lie well within the dynamical range of the FTi control loop, and thus in the linear range of the sigmoidal curve.^81^ Linear fits are thus a sufficiently precise estimation for the purposes of this study and will, at worst, underestimate any effects reported here.

In Figure 3, only data from linear fits whose R^2^ value is equal or greater than |0.6| were included. In all figures, N refers to the number of animals, n to the number of stimulus runs. For each recording, the data of no more than one stimulus run of increasing amplitude and velocity, respectively, were included in the analysis. Significance was tested with paired t-tests, significance levels: * 0.05, ** 0.01, *** 0.001. All analyses were done in Matlab (R2020a, Mathworks, RRID:SCR_001622). Figures were created using Adobe Illustrator CS6 (v16, Adobe Systems, RRID:SCR_010279).

